# Class IIa HDACs inhibit cell death pathways and protect muscle integrity in response to lipotoxicity

**DOI:** 10.1101/2022.10.17.512492

**Authors:** Sheree D. Martin, Timothy Connor, Andrew Sanigorski, Kevin A. McEwen, Darren C. Henstridge, Brunda Nijagal, David De Souza, Dedreia L. Tull, Peter J. Meikle, Greg M. Kowalski, Clinton R. Bruce, Paul Gregorevic, Mark A. Febbraio, Fiona M. Collier, Ken R. Walder, Sean L. McGee

## Abstract

Lipotoxicity, the accumulation of lipids in non-adipose tissues, alters the metabolic transcriptome and mitochondrial metabolism in skeletal muscle. The mechanisms involved remain poorly understood. Here we show that lipotoxicity increased histone deacetylase 4 (HDAC4) and histone deacetylase 5 (HDAC5), which reduced the expression of metabolic genes and oxidative metabolism in skeletal muscle, resulting in increased non-oxidative glucose metabolism. This metabolic reprogramming was also associated with impaired apoptosis and ferroptosis responses, and preserved muscle cell viability in response to lipotoxicity. Mechanistically, increased HDAC4 and 5 decreased acetylation of p53 at K120, a modification required for transcriptional activation of apoptosis. Redox drivers of ferroptosis derived from oxidative metabolism were also reduced. The relevance of this pathway was demonstrated by overexpression of loss-of-function HDAC4 and HDAC5 mutants in skeletal muscle of obese *db/db* mice, which enhanced oxidative metabolic capacity, increased apoptosis and ferroptosis and reduced muscle mass. This study identifies HDAC4 and HDAC5 as repressors of skeletal muscle oxidative metabolism, which is linked to inhibition of cell death pathways and preservation of muscle integrity in response to lipotoxicity.

## INTRODUCTION

Obesity is driving an epidemic of chronic disease, including type 2 diabetes. Linked to these diseases are adverse cellular responses arising from the accumulation of lipids in non- adipose tissues, a process termed lipotoxicity^1^. The deleterious effects of lipotoxicity include altered cellular signalling, metabolic reprogramming and cell death^2^.

Lipotoxicity alters skeletal muscle metabolism, inducing insulin resistance and impairments in aspects of oxidative metabolism. Reduced rates of both glucose and lipid oxidation have been observed in the skeletal muscle of obese subjects^3, 4^ and data describing impairments in amino acid metabolism are also emerging^5,6,7^. Experimental induction of acute lipotoxicity in humans through lipid infusion results in many of the same skeletal muscle metabolic alterations^8, 9^. These impairments in oxidative metabolism are associated with reduced expression of a metabolic and mitochondrial transcriptional program controlled by the peroxisome proliferator-activated receptor-gamma coactivator 1 alpha (PGC-1α) transcriptional coactivator^10^, encoded by the *PPARGC1A gene* . It is not known why repression of the metabolic transcriptome and reduced oxidative metabolism is an adaptation to lipotoxicity and the molecular pathways involved remain unclear.

Studies from our group and others have found that the class IIa histone deacetylase (HDAC) isoforms 4 and 5 are regulators of PGC-1α expression in skeletal muscle^11, 12^. HDAC4 and 5 act co-operatively as heterodimers to repress the myocyte-enhancer factor 2 (MEF2) family of transcription factors^13, 14^. Binding sites for the MEF2 transcription factor are found in the *PPARGC1A* promoter^12^ and in the promoters and enhancer regions of other genes involved in oxidative metabolism^11^. However, the role of the class IIa HDAC transcriptional repressors in the context of lipotoxicity is unknown. We hypothesised that HDAC4 and 5 are increased in response to lipotoxicity and contribute to the suppression of metabolic genes and oxidative metabolism and sought to understand why these adaptations occur.

## MATERIALS AND METHODS

### Cell culture

Mouse C2C12 myoblasts and myotubes were cultured using standard methods. Cells stably overexpressing HDAC4 and 5 were analysed as myoblasts as HDAC4 and 5 prevented myogenic differentiation of these cells to myotubes. Treatment doses of palmitate, camptothecin, Erastin, 1S,3R-RSL 3 (Merck, Melbourne, Australia) and Ferrostatin (Selleckchem, Houston, Tx, USA) are reported in Figure legends and were applied for 24h unless otherwise stated. Palmitate was dissolved in ethanol at 70°C and diluted in ultrapure H_2_O before conjugation to fatty acid-free BSA at 1:10 ratio (w/v). An ethanol/BSA solution without palmitate was prepared at the same time for use as a vehicle.

### Animal models

All animal experimentation was approved by the Deakin University Animal Ethics Committee (G27-2012 and G21-2015). Mice were acquired from the Animal Resources Centre (Perth, Australia) and were group housed with a 12hr light/dark cycle, temperature 21± 3°C, humidity 30-70% with *ad libitum* access to standard rodent chow and water. Heterozygous *db/+* and homozygous *db/db* mice on a C57BL/6J background were fasted for 5 hr prior to humane killing by cervical dislocation for rapid excision of tissues, which were snap frozen in liquid nitrogen. C57BL/6J mice were administered recombinant serotype 6 adeno-associated viral (AAV6) (rAAV6) vectors via intramuscular injection into the anterior and posterior compartments of the lower hind limbs. Each compartment of the left limb received 2e^10^ vector genomes (vg) of empty rAAV6, whilst each compartment of the right limb received 1e^10^ vg of WT HDAC4 rAAV6 and 1e^10^ vg of WT HDAC5 rAAV6. *db/db* mice were similarly administered rAAV6 vectors via intramuscular injection. Each compartment of the left limb received 5e^10^ vg of empty rAAV6, whilst each compartment of the right limb received 2.5e^10^ vg of DN HDAC4 rAAV6 and 2.5e^10^ vg of DN HDAC5 rAAV6. Experiments were performed ∼8 weeks after rAAV6 administration. At the conclusion of experiments mice were fasted for 4 hours before humane killing by cervical dislocation. Tissues were rapidly excised and snap frozen in liquid nitrogen or fixed in 10% formalin.

### Respiration and mitochondrial function analysis

Mitochondrial function in cells treated with vehicle (BSA) or palmitate was assessed as previously described^15^ and respiratory analysis of skeletal muscle biopsies was also performed as previously described^11^.

### Lipidomics and metabolite analysis

To determine cellular lipid profiles, ∼1e^6^ cells were collected in 100µl of PBS and frozen at - 80°C, or ∼10mg of muscle was homogenised in 100µl of PBS, and lipids were determined as previously described ^16^. For detection of Glu and GSH, cells in 6-well plates underwent metabolic arrest by washing in 1mL ultrapure water before being frozen instantaneously by the addition of liquid nitrogen. To extract polar metabolites, ice-cold methanol:chloroform (9:1) containing internal standards (0.5nM 13C-sorbitol and 5nM 13C, 15N-valine) was added. Metabolites were detected using LC and high resolution QTOF mass spectrometry previously described^17^ with the following modifications. Samples (10μL) were injected onto an Agilent 1290 LC fitted with a SeQuant ZIC®-pHILIC column (2.1 x 150 mm, 5μm; Merck) using 20mM ammonium carbonate, pH 9.0 (Sigma-Aldrich) and 100% acetonitrile as the mobile phases.

### Microarray and gene expression analysis

For comparative transcriptomics analysis, each group consisted of three biological replicates and microarray analysis was performed as previously described^11^ and conform to MIAME guidelines. Genes found to be significantly altered in the same direction in both cell and skeletal muscle data sets (n = 236) were analysed by gene set enrichment analysis using KEGG pathways. Gene expression was measured using real time RT-PCR as previously described^18^. Primer sequences are available in Table S1.

### Immunoprecipitation and western blot analysis

Cells were collected in lysis buffer (50mM Tris pH 7.5, 1mM EDTA, 1mM EGTA, 10% glycerol, 1% Triton X-100, 50mM NaF, 5mM sodium pyrophosphate, 1mM DTT, 1X protease inhibitor cocktail (Merck)) and ∼20mg skeletal muscle was homogenised in 10 volumes of lysis buffer using a handheld homogeniser. Protein concentration was determined using the BCA assay (Pierce). Immunoprecipitation and western blotting were performed as previously described^19^. Antibody details are listed in Table S2.

### Insulin/isotopic glucose administration and stable isotope metabolomics

Mice were administered 200uCi/kg of 2[1,2-^3^H]-deoxyglucose and 100uCi/kg of 1-^14^C glucose, with or without 1.2U/kg insulin, following a 5hr fast. Blood glucose was determined using a handheld glucometer (Accu-Chek Performa, Merck) and ∼5μL blood was obtained from the tail tip prior to tracer administration and 5, 15 and 30min after tracer administration. Plasma tracer concentration and tissue 2-deoxyglucose clearance were determined as we have previously described ^20^. To determine 1-^14^C glucose incorporation into glycogen, ∼10-15mg of tibialis anterior muscle was digested in 1M KOH at 70°C for 20 min and glycogen was precipitated with saturated Na_2_SO_4_, washed twice with 95% ethanol and resuspended in acetate buffer (0.84% sodium acetate, 0.46% acetic acid, pH 4.75) containing 0.3mg/mL amyloglucosidase (Merck). Glycogen was digested overnight at 37°C before glucose content was quantified using the glucose oxidase method^15^ and ^14^C-glucose incorporation was measured. To determine 1-^14^C glucose incorporation into lipids, 5-10mg of TA muscle was homogenised in chloroform/methanol (2:1) and mixed overnight at room temperature. Organic and inorganic phases were separated, and the lower organic phase was collected and evaporated under N at 45°C. The dried extract was resuspended in absolute ethanol and TG content was assayed using TG GPO-PAP reagent (Merck) and ^14^C- glucose incorporation was measured.

To assess oxidative glucose utilisation, mice were administered a bolus of 50mg of [U-^13^C] glucose (Sigma-Aldrich) via oral gavage. At 60 min later, mice were humanely killed by cervical dislocation and skeletal muscles were immediately collected. Targeted metabolomics and analysis of [U-^13^C] labelling was performed as we have previously described^20^.

### Measurement of apoptosis and cell viability

Analysis of apoptotic cells was performed by dual staining with FITC-conjugated Annexin V (Merck) and propidium iodide (PI; Thermo-Fischer, Waltham, MA, USA) and was optimised for adherent cells. For measurement of apoptosis in tissues, caspase 3 activity was measured using Caspase-Glo 3/7 assay (Promega, Alexandria, Australia). Quantification of viable cells was performed by staining with 7-aminoactinomycin D (7-AAD; Merck) and flow cytometry or with crystal violet staining with colorimetric detection.

### Reactive oxygen species detection

In cells, hydrogen peroxide (H_2_O_2_) was detected using Amplex Red assay (Invitrogen, Mt Waverly, Australia). Cells were seeded into a 96-well plate at 25,000 cells/well. The following day, cells were treated with vehicle (DMSO), 100µM mitoTEMPO or 100µM apocynin for 30 min at 37°C. Cells were then co-incubated with 50µM Amplex Red and 0.1U/ml horseradish peroxidase (HRP) for a further 30 min at 37°C. Mitochondrial ROS was designated as that sensitive to MitoTEMPO, while NADPH oxidase ROS was designated as that sensitive to apocynin. For tissues, the OxiSelect In Vitro ROS/RNS Assay kit (Green Fluorescence) (Cell Biolabs Inc., San Diego, CA, USA) was used according to the manufacturer’s instructions.

### GPX4 activity and lipid peroxidation

Glutathione dependent peroxidase activity was measured using a colorimetric Glutathione Peroxidase Assay Kit (AbCam). Cells were plated into 10cm tissue culture plates at 4x10^6^ cells per dish. After 3hrs, media was refreshed and cells incubated for 1hr, washed and collected in 250ul of supplied assay buffer. For tissues, muscle lysate was analysed. Data was then analysed as described by the manufacturer with results normalized to protein concentration. Lipid peroxidation in both cells and muscle lysates was assessed using a lipid peroxidation kit (Sigma). The assay was performed according to manufacturer’s instructions, including addition of butanol to the reaction.

### Histology

Fixed TA muscles were embedded in paraffin before transverse sections were obtained at 5μm thickness and stained for H&E and Masson’s trichrome. Slides were scanned (Aperio, Leica Biosystems) and were analysed using photoshop and image J.

### Statistical analysis

All data are expressed as mean ± SEM. Individual data points identified as greater than two standard deviations away from the mean were designated as outliers and removed. Data was assessed for normality using the Shapiro-Wilk test. Differences between groups were assessed using t-test, Kruskal-Wallis test, one-way ANOVA or two-way ANOVA as appropriate using GraphPad Prism. Specific differences between groups were identified using Dunn’s or Tukey’s multiple comparisons tests. For animal experiments where opposing hind limbs were administered different rAAVs, such that control and experimental conditions were contained within the one animal, paired t-tests were used. Differences were considered statistically significant where p<0.05.

## RESULTS

### HDAC4 and 5 are increased in models of lipotoxicity

To test the hypothesis that HDAC4 and 5 are increased by lipotoxicity, C2C12 myotubes were exposed to 0.25 or 0.5mM palmitate, or BSA (vehicle), for 16 hr. Application of 0.5mM palmitate induced alterations consistent with lipotoxicity in muscle cells, including increased ceramide, diglyceride (DG) and triglyceride (TG) levels (Fig. 1A and S1A), lower *Ppargc1a* gene expression (Fig. 1B) and a reduction in cellular respiration (Fig. 1C). The abundance of both HDAC4 and HDAC5 protein was increased by 0.5mM palmitate, while HDAC5 protein abundance was also increased by 0.25mM palmitate (Fig. 1D). There were no differences in *Hdac4* or *Hdac5* gene expression between groups (Fig. S1B).

**Figure 1:**
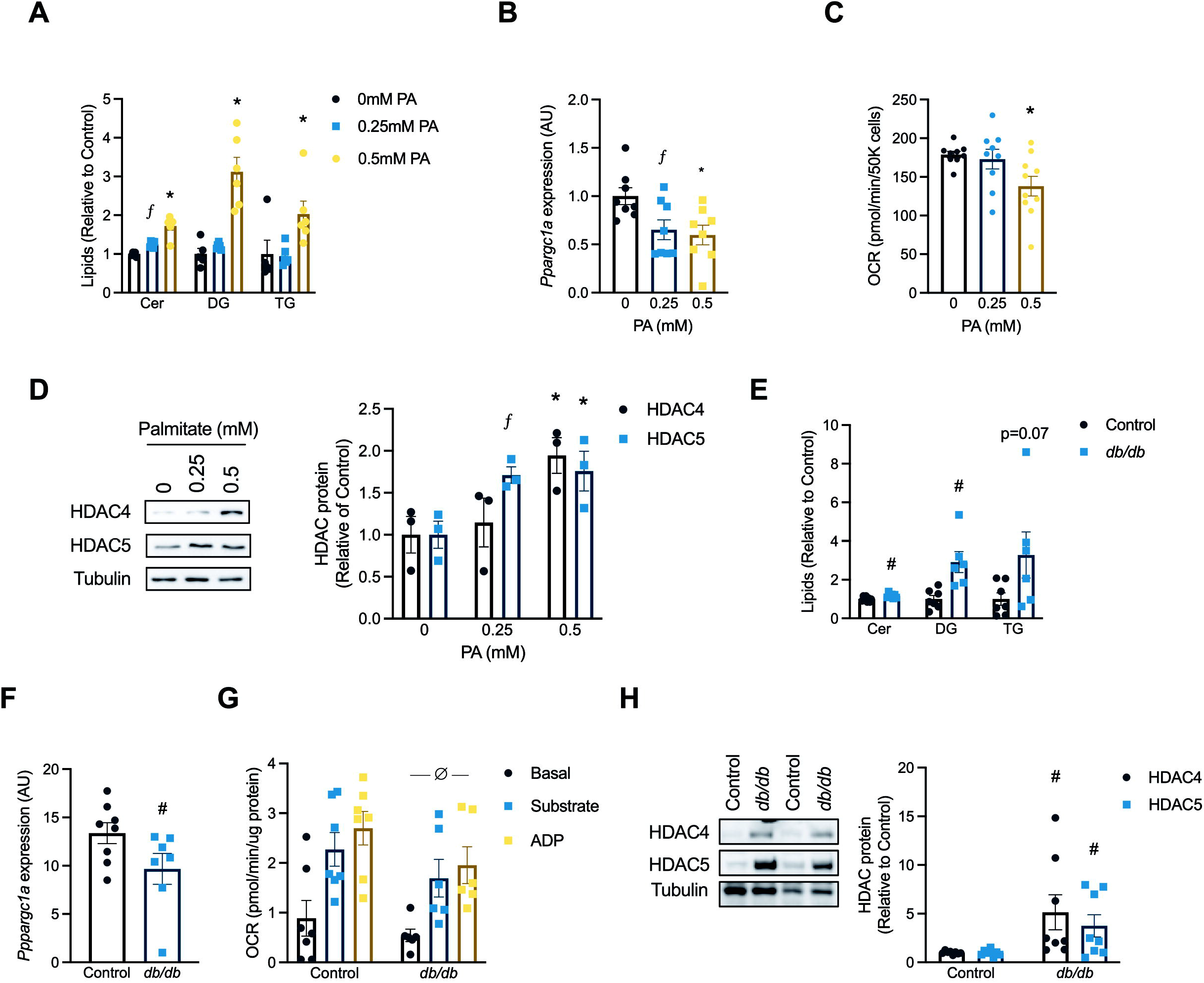
HDAC4 and 5 are increased in models of lipotoxicity. (A) Total Cer, DG and TG lipids (One-way ANOVA; Cer *p*<0.0001 (F(2,14)=73.86, ^ƒ^Tukey’s multiple comparisons test *p*<0.0077 vs. 0mM, *Tukey’s multiple comparisons test *p*<0.0001 vs. 0mM and vs 0.25mM; DG *p*<0.0001 (F(2,14)=38.50 *Tukey’s multiple comparisons test *p*<0.0001 vs. 0mM and vs 0.25mM; TG *p*<0.0017 (F(2,14)=10.38 *Tukey’s multiple comparisons test *p*=0.0333 vs. 0mM); (B) *Ppargc1a* gene expression (One-way ANOVA *p*<0.0161 (F(2,21)=5.054, ^ƒ^Tukey’s multiple comparisons test *p*<0.0484 vs. 0mM *Tukey’s multiple comparisons test *p*=0.0210 vs. 0mM); (C) Oxygen consumption rate (OCR; One-way ANOVA *p*<0.0255 (F(2,25)=4.262 *Tukey’s multiple comparisons test *p*<0.0001 vs. 0mM and vs 0.25mM), and; (D) HDAC4 and 5 protein in C2C12 myotubes treated with 0mM (BSA vehicle), 0.25mM or 0.5mM palmitate (PA) for 16 hrs (HDAC4, Kruskal-Wallis test, *p*=0.049 (*X*^2^ = 5.591), *Dunn’s multiple comparison test *p*=0.0380; HDAC5, One-way ANOVA, *p*=0.038 (F(2,6)=5.903) ^ƒ^Tukey’s multiple comparisons test *p*=0.0318, *Tukey’s multiple comparisons test *p*=0.0251). (E) Total Cer, DG and TG lipids (Unpaired t-tests, Cer *p*=0.0094, DG *p*=0.0022, TG *p*=0.0616); (F) *Ppargc1a* gene expression (Unpaired t-test, *p*=0.0357); (G) Basal, substrate and ADP-driven OCR from muscle biopsies (Two-way ANOVA, ^∅^genotype *p*=0.0493, *F*(1,33)=4.166), and; (H) HDAC4 and 5 protein in Control and *db/db* mice (Unpaired t-tests, ^#^HDAC4 *p*=0.0375, ^#^HDAC5 *p*=0.0308). Data are mean±SEM, n=3-6 biological replicates per group for cell experiments and 6-8 per group for animal experiments.

To examine whether skeletal muscle HDAC4 and 5 are increased in an *in vivo* model of lipotoxicity, the TA skeletal muscle of obese *db/db* and control heterozygous littermate mice was analysed. Total ceramide and DG were were increased in skeletal muscle of *db/db* mice, while there was a trend for TG to also be higher (Fig. 1E and S1C). Increased muscle lipid concentrations in were associated with decreased *Ppargc-1α* gene expression (Fig. 1F), reduced skeletal muscle respiratory responses (Fig. 1G) and increased HDAC4 and 5 protein abundance (Fig. 1H). Skeletal muscle gene expression levels of *Hdac5*, but not *Hdac4*, were increased in *db/db* mice (Fig. S1D). Notwithstanding the complex physiology of the *db/db* model, these data suggest that skeletal muscle HDAC4 and 5 are increased in response to lipotoxicity and are associated with reduced *Ppargc1a* expression and impaired oxidative metabolism.

### Overexpression of HDAC4 and 5 in skeletal muscle represses Pppargc1a expression and oxidative capacity

To examine whether increased HDAC4 and 5 impacts skeletal muscle metabolism, a bilateral skeletal muscle HDAC4 and 5 mouse model was developed using rAAV6 vectors expressing HDAC4 and HDAC5 (Fig. 2A). In TA muscle, this model had increased *Hdac4* and *Hdac5* gene expression (Fig. 2B) and protein (Fig. 2C). Overexpression of HDAC4 and 5 reduced *Ppargc1a* expression (Fig. 2D) and the expression of a range of MEF2 and PGC-1α target genes involved in oxidative and mitochondrial metabolism (Fig. 2E). These changes in metabolic transcripts were associated with reduced protein abundance of ATP synthase subunits, but not other subunits of electron transport chain complexes (Fig. 2F). Reduced ATP synthase components was associated with blunted respiratory responses in muscles overexpressing HDAC4 and 5 when driven by both succinate (Fig. 2G) and malate (Fig. 2H) as substrates. Overexpression of HDAC4 and 5 was not associated with lipid accumulation (Fig. 2I). These data indicate that increasing HDAC4 and 5 represses *Ppargc1a* gene expression and oxidative capacity in skeletal muscle.

**Figure 2:**
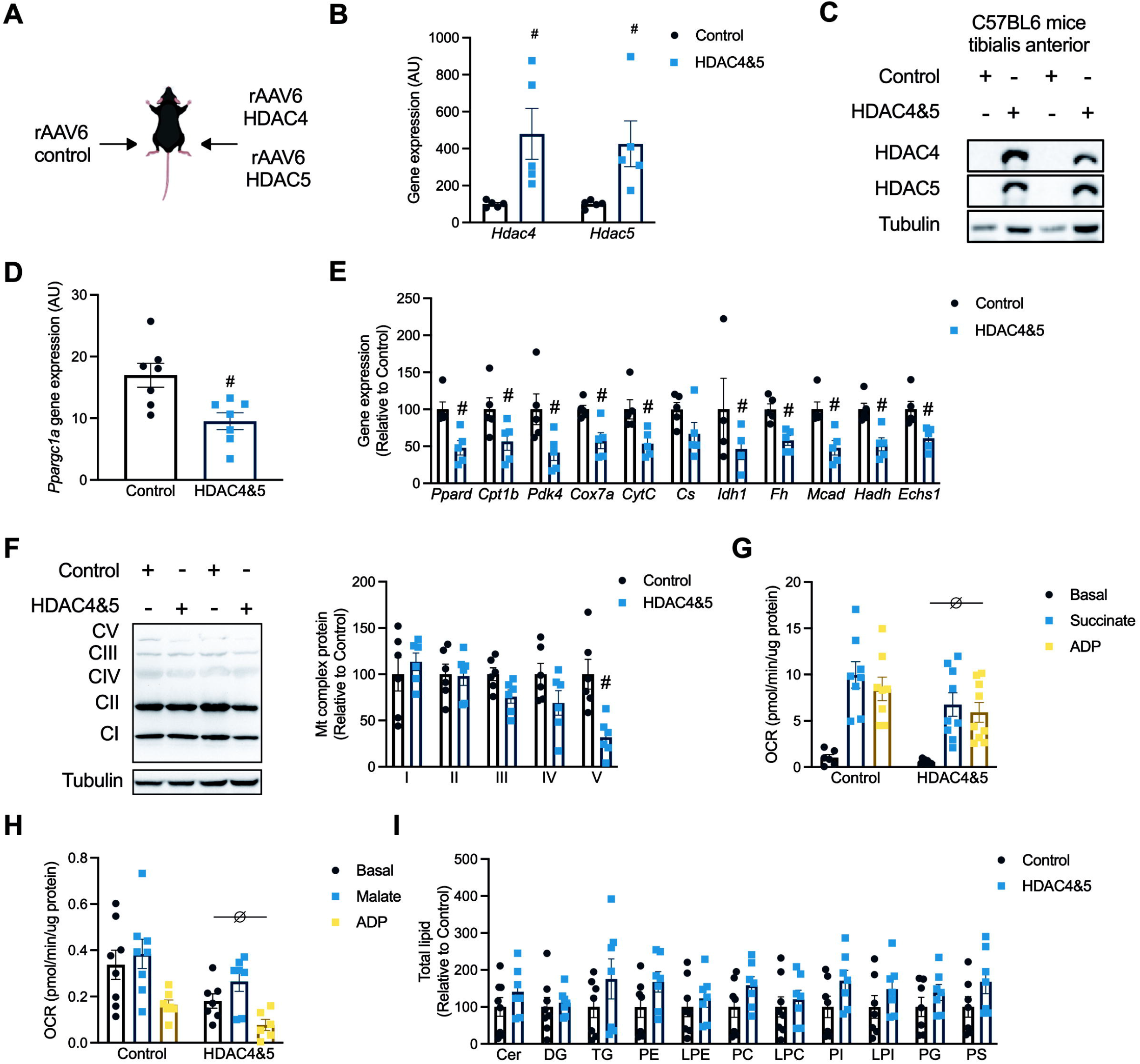
Overexpression of HDAC4 and 5 in skeletal muscle suppresses Pppargc1a expression and oxidative capacity. (A) The bilateral AAV6 HDAC4 and 5 overexpression mouse model. (B) *Hdac4* and *Hdac5* gene expression (Paired t-test, ^#^*Hdac4 p* =0.0246, ^#^*Hdac5 p* =0.0305); (C) HDAC4 and 5 protein; (D) *Ppargc1a* gene expression (Paired t-test, *p*=0.0006); (E) Metabolic gene expression (Paired t-test, ^#^*Ppard p*=0.0055, ^#^*Cpt1b p*=0.0106, ^#^*Pdk4 p* =0.0029, ^#^*Cox7a p* =0.0087, ^#^*CytC p* =0.0020, ^#^*Idh1 p* =0.0179, ^#^*Fh p* =0.0015, ^#^*Mcad p*=0.0028, ^#^*Hadh p*=0.0029, ^#^*Echs1 p*=0.0023); (F) Respiratory chain complex subunit protein (Paired t-test, ^#^Complex V *p*<0.0001); (G) Basal, succinate and ADP-driven oxygen consumption rate (OCR) from muscle biopsies (Two-way ANOVA, ^∅^genotype *p*=0.0247, *F*(1,43)=5.417); (H) Basal, malate and ADP-driven OCR from muscle biopsies (Two-way ANOVA, ^∅^genotype *p*=0.0036, *F*(1,37)=9.661), and; (I) Ceramide (Cer), diglyceride (DG), triglyceride (TG), phosphatidylethanolamine (PE), lysophosphatidylethanolamine (LPE), phosphatidylcholine (PC), lysophosphatidylcholine (LPC), phosphatidylinositol (PI), lysophosphatidylinositol (LPI), phosphoglycerol (PG) and phosphatidylserine (PS) lipids in Control and HDAC4 and 5 overexpressing skeletal muscle. Data are mean±SEM, n=4-6 per group.

### Overexpression of HDAC4 and 5 in skeletal muscle inhibits oxidative glucose utilisation

Reduced skeletal muscle oxidative capacity associated with lipotoxicity is linked with reduced insulin action and impaired glucose utilisation^4, 21, 22^. In bilateral skeletal muscle HDAC4 and 5 mice administered either vehicle or insulin and glucose tracers, glucose clearance was increased in skeletal muscles overexpressing HDAC4 and 5 (Fig. 3B). When expressed relative to control skeletal muscle within each mouse, HDAC4 and 5 overexpression increased glucose clearance in vehicle but not insulin-stimulated conditions (Fig. 3C). Insulin reduced the excursion of both glucose tracers (Fig. S2A-B). Overexpression of HDAC4 and 5 increased glucose incorporation into glycogen (Fig. 3D) and increased glycogen concentration (Fig. S2C). In contrast, insulin increased glucose incorporation into lipids (Fig. 3E), which was associated with increased Akt phosphorylation at T308 (Fig. 3G) and S473 (Fig. 3H). Consistent with glucose clearance, phosphorylation of TBC1D4 at S642 was increased in skeletal muscle overexpressing HDAC4 and 5 (Fig. 3I).

**Figure 3:**
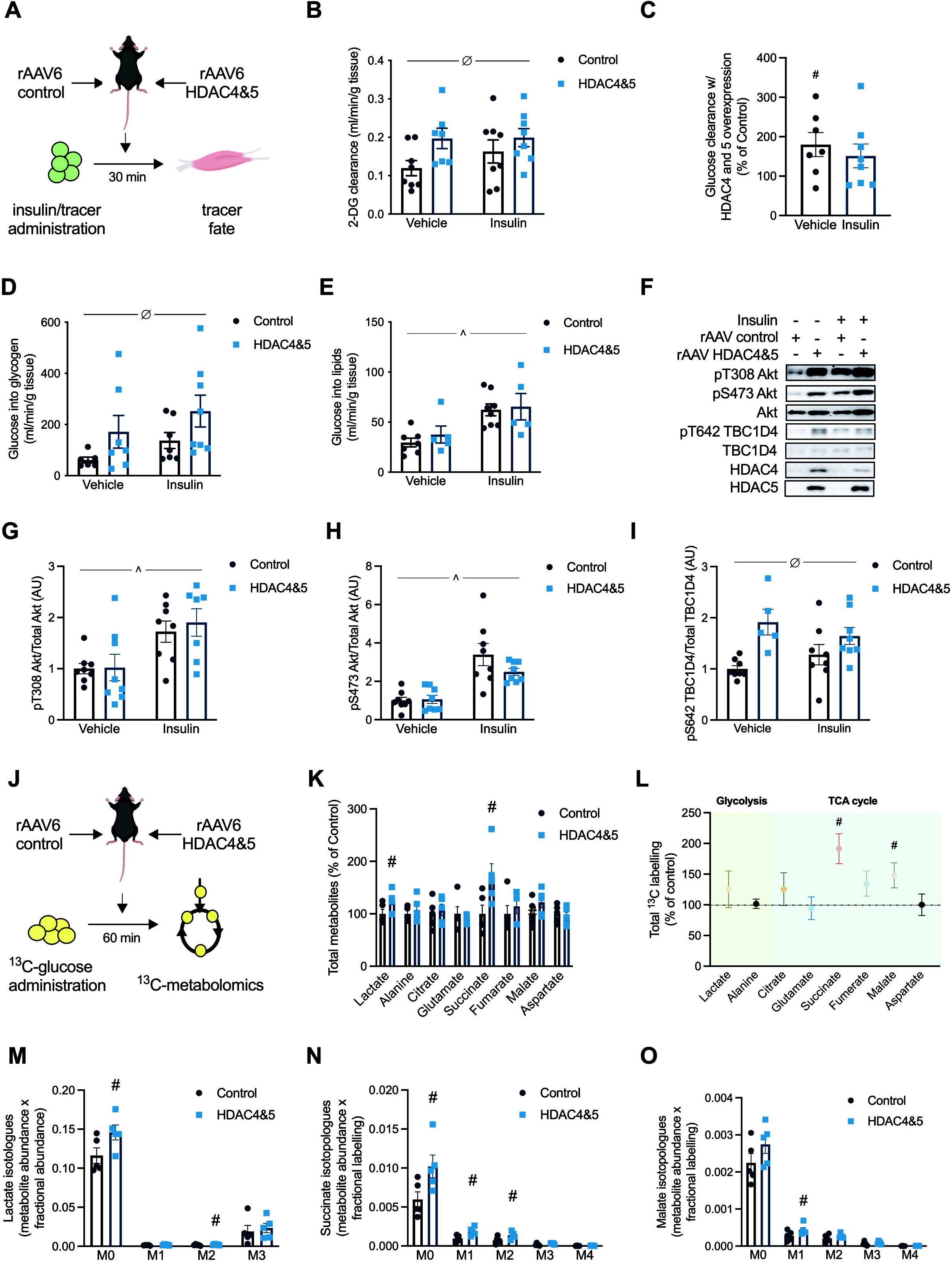
Overexpression of HDAC4 and 5 in skeletal muscle inhibits oxidative glucose utilisation. (A) Experimental overview of insulin and isotopic glucose tracers in the bilateral AAV6 HDAC4 and 5 overexpression model. (B) Glucose clearance (Two-way ANOVA, ^∅^genotype *p*=0.0328, *F*(1,27)=5.061); (C) Glucose clearance in muscle overexpressing HDAC4 and 5 relative to Control (Paired t-test, ^#^Vehicle *p*=0.0355); (D) ^14^C-glucose incorporation into glycogen (Two-way ANOVA, ^∅^genotype *p*=0.0372, *F*(1,24)=4.864); (E) ^14^C-glucose incorporation into lipid (Two-way ANOVA, ^Insulin *p*=0.0009, F(1,21)=14.94); (F) Representative images of insulin signalling components; (G) pT308 Akt (Two-way ANOVA, ^Insulin *p*=0.0010, F(1,27)=13.75); (H) pS473 Akt (Two-way ANOVA, ^Insulin *p*<0.0001, F(1,28)=33.13), and; (I) pS642 TBC1D4 (Two-way ANOVA, ^∅^genotype *p*=0.0011, *F*(1,25)=13.60) in bilateral AAV6 HDAC4 and 5 mice administered insulin and isotopic glucose tracers. (J) Experimental overview of stable isotope metabolomics in the bilateral AAV6 HDAC4 and 5 overexpression model. (K) Total metabolite abundance (Paired t-test, ^#^Lactate *p*=0.0447, ^#^Succinate *p*=0.0355); (L) Total ^13^C labelling of glycolytic and TCA cycle metabolites (Paired t-test, ^#^Succinate *p*=0.0447, ^#^Malate *p*=0.0355); (M) Lactate isotopologue (Paired t-test, ^#^M0 *p*=0.0448, ^#^M2 *p*=0.0299); (N) Succinate isotopologues (Paired t-test, ^#^M0 *p*=0.0230, ^#^M1 *p*=0.0002, ^#^M2 *p*=0.0005), and; (O) Malate isotopologues (Paired t-test, ^#^M1 *p*=0.0178) in bilateral AAV6 HDAC4 and 5 mice administered ^13^C- glucose tracer. (E) Data are mean±SEM, n=5-8 per group.

This was further explored using a targeted stable isotope metabolomics approach in bilateral skeletal muscle HDAC4 and 5 mice (Fig. 3J). Overexpression of HDAC4 and 5 increased the abundance of lactate and succinate (Fig. 3K) and total ^13^C labelling of succinate and malate (Fig. 3L). Isotopologue analysis revealed accumulation of both M+0 and M+2 lactate in muscle overexpressing HDAC4 and 5 (Fig. 3M), suggesting both increased non-oxidative glucose utilisation and increased malate/pyruvate cycle flux, respectively^23^. Succinate M+0, M+1 and M+2 isotopologues were increased in muscle overexpressing HDAC4 and 5 (Fig. 3N), consistent with a major impairment in SDH activity^24^, which is bifunctional for both the TCA cycle and the electron transport chain. These data are also supported by reduced succinate-driven respiration in muscles overexpressing HDAC4 and 5 (Fig. 2G). Similarly, HDAC4 and 5 overexpression was associated with increased malate M+0 and M+1 isotopologues (Fig. 3O). Together, these data indicate that increased HDAC4 and 5 is linked with defective oxidative glucose utilisation and increased non-oxidative glucose utilisation in skeletal muscle.

### HDAC4 and 5 enhance cell survival in response to lipotoxicity and supress the apoptosis and ferroptosis cell death pathways

To better understand why this metabolic reprogramming occurs, important genes regulated by HDAC4 and 5 in the context of lipotoxicity were identified using a comparative transcriptomics approach. Specifically, the transcriptome altered in the skeletal muscle of *db/db* mice in which HDAC4 and 5 was increased (Fig. 1J) was compared to the transcriptome altered in myogenic C2C12 myoblasts stably overexpressing HDAC4 and 5 (Fig. S3A). Overexpression of HDAC4 and 5 in these cells reduced *Ppargc1a* gene expression, suppressed OXPHOS and TCA cycle genes, and reduced oxidative capacity (Fig. S3B-D). The comparative transcriptomic analysis and subsequent pathway analysis of coregulated genes revealed the p53 signalling pathway related to cell survival as the only significantly regulated pathway (Fig. S3E).

Profiling of downstream genes in this pathway in C2C12 myoblasts overexpressing HDAC4 and 5 revealed that several pro-apoptosis genes were reduced, including *Casp3* and *9, Cycs, Pmaip1* and *Bbc3* (Fig. 4A), consistent with suppression of apoptosis. Similarly, genes involved in ferroptosis, a form of iron-mediated cell death involving lipid peroxidation ^25^, were altered in ways that indicated suppression of ferroptosis, including increased *Slc7a11* expression and reduced *Sat1* and *Alox15* expression (Fig. 4A). These data suggest that increased HDAC4 and 5 inhibit apoptotic and ferroptotic cell death pathways. Indeed, overexpression of HDAC4 and 5 in C2C12 myoblasts reduced the number of apoptotic cells following exposure to the apoptosis-inducing agent camptothecin (Fig. 4B). Similarly, cell viability was higher in cells overexpressing HDAC4 and 5 following exposure to the ferroptosis-inducing agent RSL3 (Fig. 4C). These data raised the possibility that suppression of the apoptosis and ferroptosis pathways by HDAC4 and 5 enhances cell survival in response to lipotoxicity. To test this hypothesis, cells were exposed to increasing concentrations of palmitate intended to challenge cell viability. Overexpression of HDAC4 and 5 enhanced the viability of cells exposed to both 1 and 2mM of palmitate compared with control cells and resulted in cells being largely resistant to lipotoxicity (Fig. 4D). This protection of cell viability was associated with reduced cleavage of caspase 3 and 9, the irreversible events that trigger apoptosis (Fig. 4E), as well as fewer apoptotic cells (Fig. 4F) in response to increasing concentrations of palmitate. Overexposure of western blots showed that endogenous HDAC4 and 5 were increased by these high concentrations of palmitate (Fig. S3F), similar to the lower concentrations of palmitate characterised previously (Fig. 1E). Similarly, overexpression of HDAC4 and 5 reduced lipid peroxidation, the trigger for ferroptosis induced-cell death, in response to increasing concentrations of palmitate (Fig. 4H). These data indicate that increased HDAC4 and 5 inhibit the apoptosis and ferroptosis cell death pathways and enhance cell viability in response to lipotoxicity.

**Figure 4:**
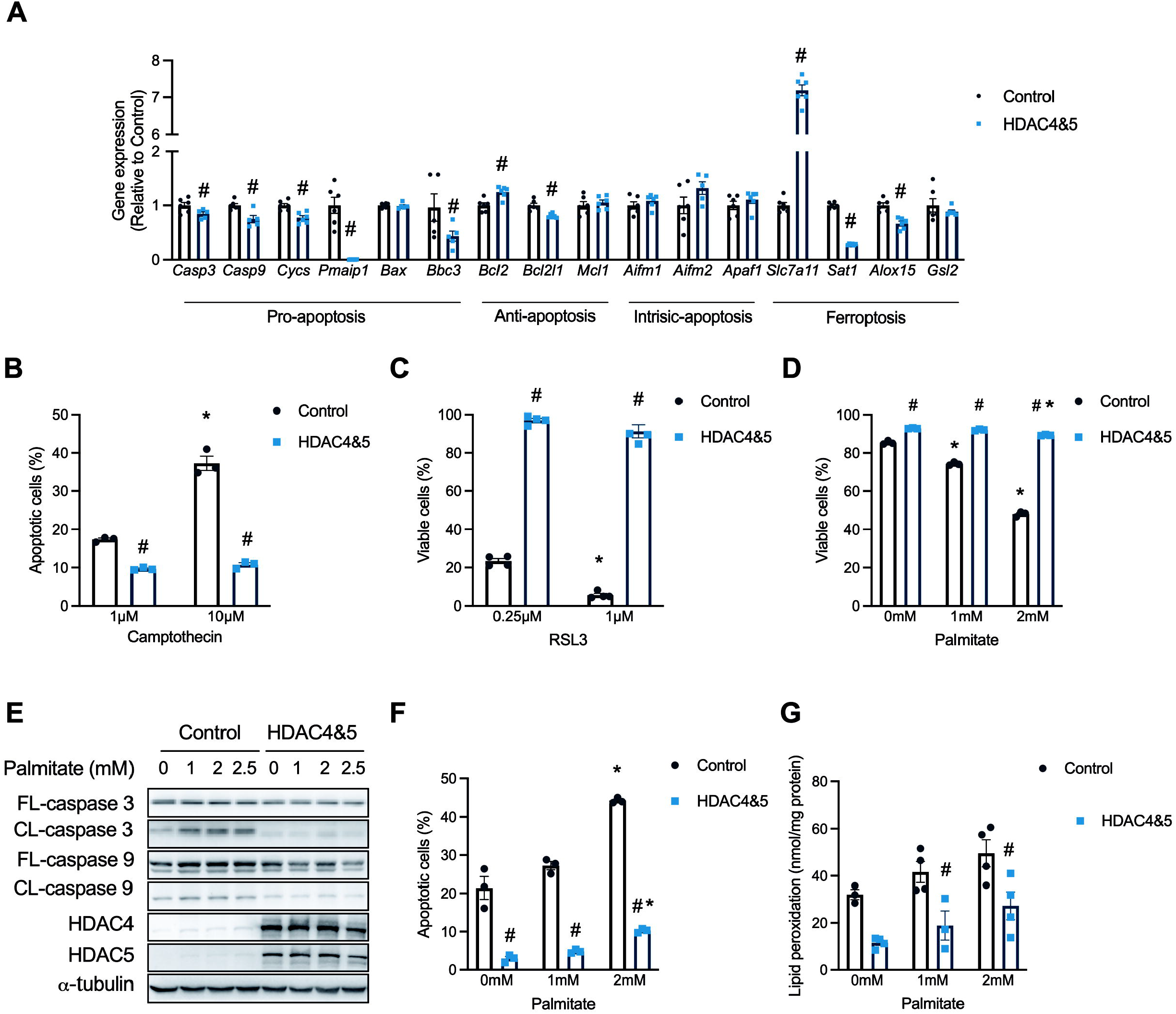
HDAC and 5 enhance cell survival in response to lipotoxicity and supress the apoptosis and ferroptosis cell death pathways. (A) Expression of p53-pathway genes involved in pro-apoptosis, anti-apoptosis, intrinsic apoptosis and ferroptosis (Unpaired t- test, ^#^*Casp3 p* =0.0451, ^#^*Casp9 p* =0.0057, ^#^*Cycs p* =0.0015, ^#^*Pmaip1 p* <0.0001, ^#^*Bbc3 p*=0.0253, ^#^*Bcl2 p* =0.0034, ^#^*Bcl2l1 p* =0.0008, ^#^*Slc7a11 p* <0.0001, ^#^*Sat1 p* <0.0001, ^#^*Alox15 p*=0.0001); (B) Apoptotic cells in response to the apoptosis-inducing agent Camptothecin (Two-way ANOVA, genotype *p*<0.0001 *F*(1,12)=349.3, treatment *p*<0.0001 *F*(2,12)=270.4, interaction *p*<0.0001 *F*(2,12)=149.5; ^#^p<0.05 vs Control for same treatment, *p<0.05 vs vehicle by Tukey’s multiple comparisons tests); (C) Cell viability in response to the ferroptosis-inducing agent RSL3 (Two-way ANOVA, genotype *p*<0.0001 *F*(1,12)=1645, treatment *p*<0.0001 *F*(1,12)=36.09, interaction *p*<0.0090 *F*(1,12)=9.665; ^#^p<0.05 vs Control for same treatment, *p<0.05 vs 0.25μM RSL by Tukey’s multiple comparisons tests); (D) Cell viability in response to palmitate (Two-way ANOVA, genotype *p*<0.0001 *F*(1,12)=3374, treatment *p*<0.0001 *F*(2,12)=1012, interaction *p*<0.0001 *F*(2,12)=684.8; ^#^p<0.05 vs Control for same treatment, *p<0.05 vs vehicle by Tukey’s multiple comparisons tests); (E) Full- length (FL) and cleaved (CL) caspase 3 and 9 in response to palmitate; (F) Apoptotic cells in response to palmitate (Two-way ANOVA, genotype *p*<0.0001 *F*(1,12)=1468, treatment *p*=0.0001 *F*(2,12)=20.35, interaction *p*<0.0050 *F*(2,12)=8.498; ^#^p<0.05 vs Control for same treatment, *p<0.05 vs vehicle by Tukey’s multiple comparisons tests); (G) Lipid peroxidation in response to palmitate (Two-way ANOVA, genotype *p*<0.0001 *F*(1,12)=313.4, treatment *p*<0.0103 *F*(2,12)=67.31; ^#^p<0.05 vs Control for same treatment by Tukey’s multiple comparisons tests) in Control and HDAC4 and 5 overexpressing C2C12 myoblasts. Data are mean±SEM, n=3-4 biological replicates per group.

### HDAC4 and 5 inhibit apoptosis by reducing p53 K120 acetylation

In several cancer cell types, HDAC5 has been linked with reduced acetylation of p53 at K120, which inhibits p53 transcriptional activity and the expression of apoptosis genes, which reduces apoptosis sensitivity^26,27,28^. Furthermore, acetylation of p53, including at K120, can contribute to the transcriptional regulation of ferroptosis^29^. To explore whether reduced acetylation of p53 at K120 is involved in suppression of apoptosis and ferroptosis by HDAC4 and 5 in myogenic cells, C2C12 myoblasts stably overexpressing HDAC4 and 5 were transfected with wild type p53, or p53 in which K120 was mutated to either glutamine (K120Q; acetylation mimetic gain-of function) or arginine (K120R; deacetylation mimetic loss-of-function) (Fig. 5A). The acetylation of p53 at K120 was reduced by the overexpression of HDAC4 and 5 (Fig. 5A). In cells overexpressing HDAC4 and 5, introduction of the p53^K120Q^ acetylation mimetic mutant restored the expression of the pro-apoptosis genes *Casp3, Casp9* and *Cycs* (Fig. 5B). However, the expression of *Slc7a11*, which opposes ferroptosis, was not influenced by expression of this mutant (Fig. 5B). This suggests that reduced acetylation of p53 at K120 by HDAC4 and 5 suppresses apoptosis, but not ferroptosis, transcriptional programs. Expression of p53^K120Q^ also restored apoptosis in HDAC4 and 5 overexpressing cells exposed to the apoptosis-inducing agent camptothecin (Fig. 5C). Although p53 is also known to regulate aspects of cellular metabolism^30^, gain and loss-of-function K120 p53 mutants had no effect on either oxidative (Fig. S4A) or glycolytic flux (Fig. S4B). To determine whether p53 K120 acetylation determines cell viability in response to lipotoxicity, cells expressing WT p53 or p53^K120R^ were exposed to 2mM palmitate. However, the p53^K120R^ mutant did not influence cell viability in response to palmitate exposure (Fig. 5D). Furthermore, the apoptosis inhibitor, Z-DED-FMK, had no effect on cell viability in response to palmitate exposure (Fig. 5E), suggesting that other mechanisms are also involved in the cell death response to lipotoxicity.

**Figure 5:**
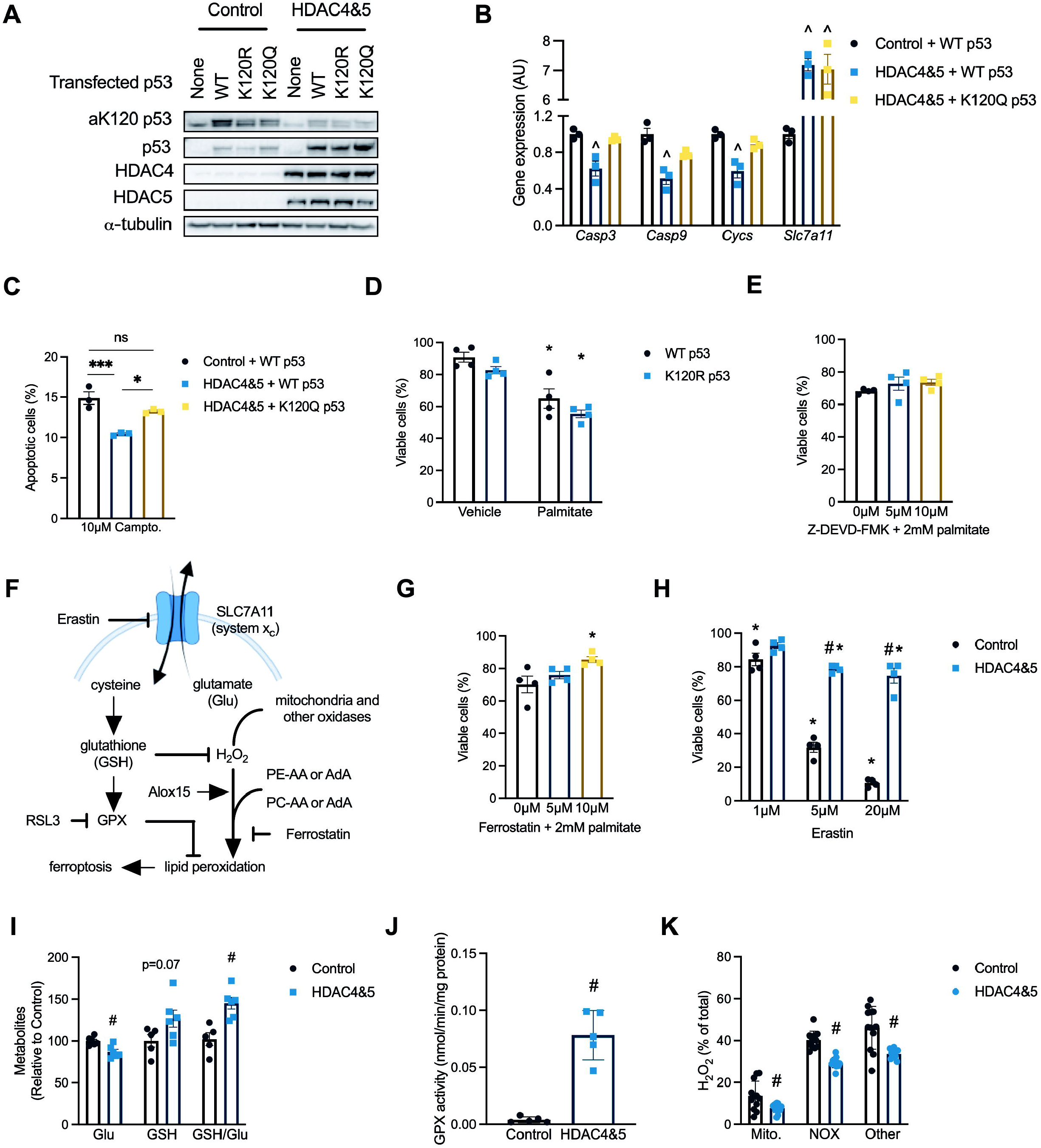
HDAC4 and 5 inhibit apoptosis by reducing p53 K120 acetylation and reduces multiple metabolic inputs to the ferroptosis pathway. (A) Acetylated p53 at lysine 120 (aK120 p53); (B) Expression of caspase 3 (*Casp3;* One-way ANOVA *p*=0.0040 *X*^2^=0.8408, ^*p*<0.05 vs Control + WT p53 and HDAC4&5 + K120Q p53 by Tukey’s multiple comparisons test), caspase 9 (*Casp9;* One-way ANOVA *p*=0.0025 *X*^2^=0.8650, ^*p*<0.05 vs Control + WT p53 and HDAC4&5 + K120Q p53 by Tukey’s multiple comparisons test), cytochrome c (*Cycs;* One- way ANOVA *p*=0.0033 *X*^2^=0.8518, ^*p*<0.05 vs Control + WT p53 by Tukey’s multiple comparisons test) and the cystine/glutamate antiporter xCT (*Slc7a11;* One-way ANOVA *p*<0.0001 *X*^2^=0.9769, ^*p*<0.05 vs Control + WT p53 by Tukey’s multiple comparisons test); (C) Percentage of apoptotic cells following exposure to camptothecin (One-way ANOVA *p*=0.0014 *X*^2^=0.8875, ^*p*<0.05 vs Control + WT p53 and HDAC4&5 + K120Q p53 by Tukey’s multiple comparisons test); (D) Cell viability 16 hrs after palmitate exposure (Two-way ANOVA, treatment *p*<0.0001 *F*(1,12)=50.44, genotype *p*=0.0361 *F*(1,12)=5.56; *p<0.05 vs vehicle for same genotype by Tukey’s multiple comparisons tests) in C2C12 myoblasts expressing wild type (WT) or K120R (loss-of-function) p53 treated with vehicle or 2mM palmitate for 24 hrs. (E) Cell viability following palmitate exposure and co-incubation with vehicle (DMSO) or increasing concentrations of Z-DEVD-FMK (One-way ANOVA *p*=0.3282); (F) Schematic showing key regulatory points in the ferroptosis pathway; (G) Cell viability following palmitate exposure and co-incubation with vehicle (DMSO) or increasing concentrations of Ferrostatin (One-way ANOVA *p*=0.0323 *X*^2^=0.5337, \**p*<0.05 vs Vehicle by Tukey’s multiple comparisons test); (H) Cell viability following exposure to increasing concentrations of Erastin (Two-way ANOVA, genotype *p*<0.0001 *F*(1,18)=293.7, treatment *p*<0.0001 *F*(2,18)=139.6, interaction *p*<0.0001 *F*(2,18)=50.88; ^#^p<0.05 vs Control for same treatment, *p<0.05 vs Vehicle same genotype by Tukey’s multiple comparisons tests); (I) Glutamate (Glu), glutathione (GSH) and the GSH/Glu ratio (Unpaired t-tests, ^#^Glu *p*=0.0049, ^#^GSH/Glu *p*=0.0023); (J) Glutathione peroxidase (GPX) activity (Unpaired t-test, *p*<0.0001), and; (K) H_2_O_2_ derived from mitochondria (mito), NADPH oxidases (NOX) and other sources (Unpaired t-tests, ^#^Mito *p*=0.0105, ^#^NAPDH Ox. *p*<0.0001, ^#^Other *p*=0.0004) in Control or HDAC4 and 5 overexpressing C2C12 myoblasts. Data are mean±SEM, n=4-10 biological replicates per group.

### HDAC4 and 5 inhibit multiple metabolic inputs to the ferroptosis pathway

The role of ferroptosis-mediated mechanisms of cell death (Fig. 5F) were further explored. The ferroptosis inhibitor Ferrostatin enhanced C2C12 myoblast viability following exposure to 2mM palmitate (Fig. 5G). Overexpression of HDAC4 and 5 enhanced cell viability in response to increasing concentrations of Erastin (Fig. 5H), an agent that induces ferroptosis by inhibiting system x and glutamate and cysteine exchange^31^, which in turn reduces glutathione (GSH) levels and results in lipid peroxidation through multiple mechanisms (Fig. 5F). Consistent with HDAC4 and 5 increasing the expression of *Slc7a11* (Fig. 4A), overexpression of HDAC4 and 5 reduced intracellular glutamate concentrations, tended to increase intracellular GSH and increased the GSH/glutamate ratio (Fig. 5I). This was associated with increased glutathione peroxidase (GPX) activity in cells overexpressing HDAC4 and 5 (Fig. 5J) and reduced total H_2_O_2_ production (Fig. S4C), including from mitochondria and NADPH oxidases (Fig. 5K). There were no differences in the levels of phosphatidylcholine (PC), lysophosphatidylcholine (LPC), lysophosphatidyl-ethanolamine (LPE) and phosphatidylethanolamine (PE) lipids that undergo peroxidation during ferroptosis^32^ in skeletal muscle of bilateral HDAC4 and 5 mice (Fig. S4D). Therefore, these data indicate that increased HDAC4 and 5 alters redox inputs to ferroptosis that are regulated by oxidative capacity.

### HDAC4 and 5 are required to suppress apoptosis and ferroptosis and maintain muscle mass in obese mice

To establish whether HDAC4 and 5 suppress apoptosis and ferroptosis in skeletal muscle in response to lipoxicity *in vivo*, a bilateral skeletal muscle loss-of-function HDAC4 and 5 model was developed in obese *db/db* mice using AAV6 vectors (Fig. 6A). This loss-of-function approach employed active-site HDAC4 and 5 mutants (D832N HDAC4 and D861N HDAC5), which we have previously described as acting in a dominant negative (DN) manner when overexpressed^11^. Overexpression of these mutants in skeletal muscle increases the expression of *Ppargc1a* and other metabolic genes and enhance oxidative metabolism^11^. Expression of DN HDAC4 and 5 to *db/db* mice increased HDAC4 and 5 gene (Fig. 6B) and protein (Fig. 6C) expression and increased the expression of *Ppargc1a* (Fig. 6D). Consistent with these findings, respiratory responses were increased in muscle expressing DN HDAC4 and 5 (Fig. 6E). Expression of DN HDAC4 and 5 increased caspase 3 activity (Fig. 6F) and lipid peroxidation (Fig. 6G), functional readouts of apoptosis and ferroptosis, respectively. Examination of the mechanisms involved in HDAC4 and 5-mediated apoptosis revealed that p53 acetylation of K120 tended to be higher (Fig. 6H), and the expression of p53-dependent pro-apoptosis genes, including *Casp3, Casp9* and *Bbc3* was increased (Fig. 6I). The ferroptosis mechanisms regulated by HDAC4 and 5 were also examined. In contrast to observations in myogenic cells, the expression of *Slc7a11* in skeletal muscle was low and close to assay detection limits (Fig. S5A). This suggests that system x_c_ likely has only a minor role in the regulation of ferroptosis in skeletal muscle. There were no differences in GPX activity between groups (Fig. S5B). However, expression of DN HDAC4 and 5 increased ROS (Fig. 6J) and increased *Sat1* and *Alox15* expression (Fig. 6K), indicating that increased capacity for lipid peroxidation. Increased apoptosis and ferroptosis in skeletal muscles expressing DN HDAC4 and 5 was associated with reduced mass of the gastrocnemius and the TA muscles (Fig. 6L). In the TA muscle, there was no difference in fibre diameter between groups (Fig. S5C). However, Masson’s trichrome staining showed sporadic areas of localised fibrosis in response to DN HDAC4 and 5 overexpression (Fig. 6M). In skeletal muscles overexpressing DN HDAC4 and 5 there were also discreet regions of fibres with centralised nuclei (Fig. 6N), indicative of regenerating muscle fibres. These data show that HDAC4 and 5 are required to suppress apoptosis and ferroptosis and to maintain muscle mass and integrity in obese mice.

**Figure 6:**
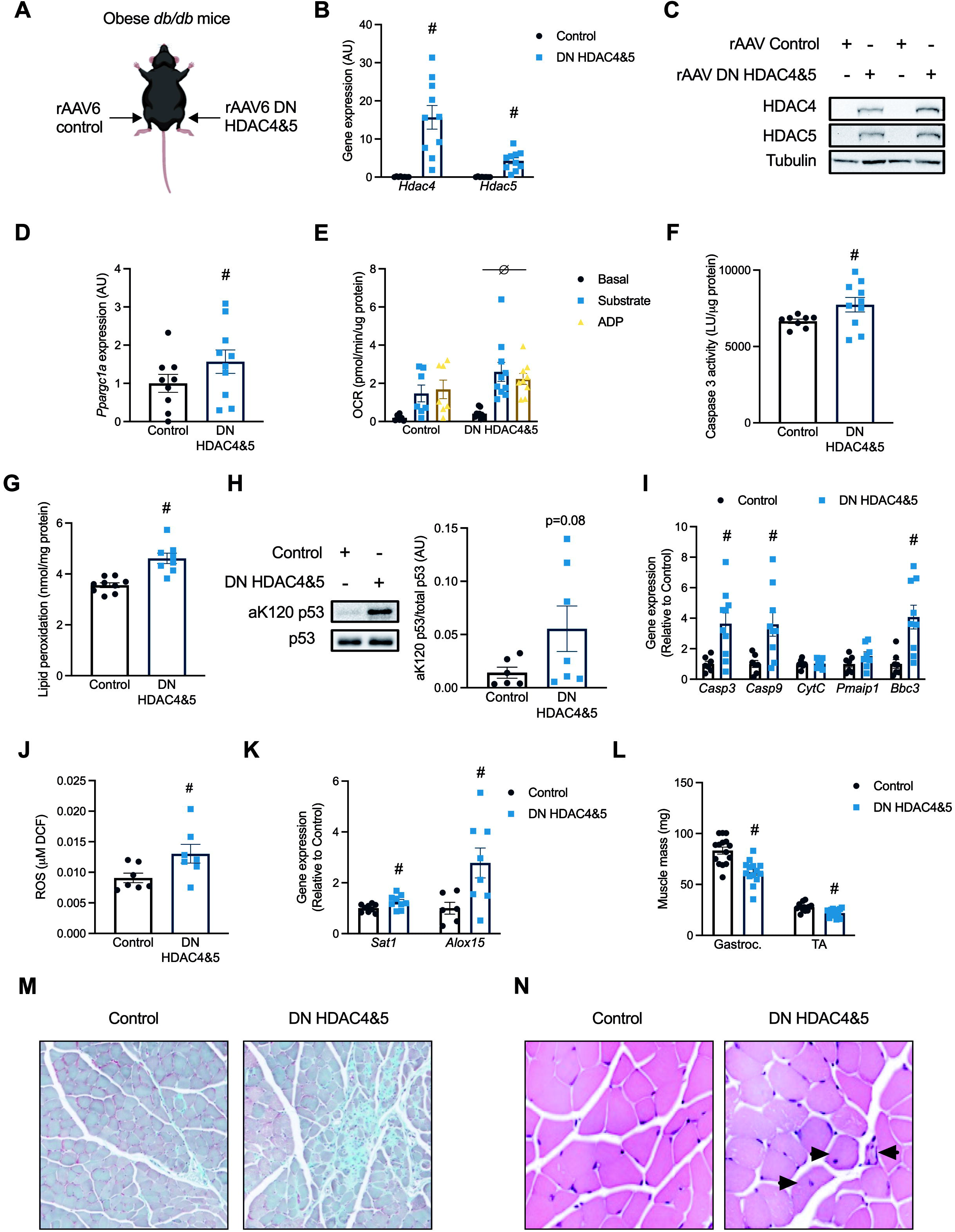
HDAC4 and 5 are required to suppress apoptosis and ferroptosis and maintain muscle mass in obese mice. (A) The bilateral AAV6 dominant negative (DN) HDAC4 and 5 overexpression *db/db* mouse model. (B) *Hdac4* and *Hdac5* gene expression (Paired t-tests, ^#^*Hdac4 p*=0.0003, ^#^*Hdac5 p*=0.0002); (C) HDAC4 and 5 protein; (D) *Ppargc1a* gene expression (Paired t-tests, *p*=0.0191); (E) Basal, substrate and ADP-driven oxygen consumption rate (OCR) from muscle biopsies (Two-way ANOVA, genotype *p*=0.0482 *F*(1,42)=4.141); (F) Caspase 3 activity (Paired t-tests, *p*=0.0292); (G) Lipid peroxidation (Paired t-tests, *p*=0.0013); (H) Acetylated p53 at lysine 120 (aK120 p53); (I) Expression of apoptosis genes (Paired t-tests, ^#^*Casp3 p*=0.0036, ^#^*Casp9 p*=0.0064, ^#^*Bbc3 p*=0.0024); (J) Reactive oxygen species (ROS; Paired t-tests, *p*=0.0292); (K) Expression of ferroptosis genes (Paired t-tests, ^#^*Sat1 p*=0.0257, ^#^*Casp9 p*=0.0286); (L) Mass of the gastrocnemius (Gastroc.) and tibialis anterior (TA) skeletal muscles (Paired t-tests, ^#^Gastroc. *P*<0.0001, ^#^TA *p*=0.0004); (M) Masson’s trichrome staining, and; (N) H&E staining of sections of the TA muscle in Control and DN HDAC4 and 5 overexpressing skeletal muscle in *db/db* mice. Arrows indicate fibres with centralised nuclei. Data are mean±SEM, n=6-10 per group.

## DISCUSSION

Lipotoxicity has a major impact on the metabolism, function and survival of various tissues including skeletal muscle. The current study has revealed that increases in HDAC4 and 5 in response to lipotoxicity induce transcriptional repression of oxidative metabolism and of the apoptosis and ferroptosis cell death pathways in both myogenic cells and skeletal muscle. The impairment in oxidative metabolism also reduces redox inputs to the ferroptosis pathway. These data indicate that metabolic reprogramming in skeletal muscle in response to lipotoxicity is directly linked with cell survival responses.

Previous findings revealed that lipotoxicity is associated with suppression of a pro-apoptosis transcriptional program in skeletal muscle, with consequences for maintenance of muscle mass^33^. Our data show that the class IIa HDACs are essential for this response, as loss of HDAC4 and 5 function in skeletal muscle in a model of lipotoxicity increased apoptosis genes and markers of apoptosis, which was associated with reduced muscle mass. Furthermore, a readout of ferroptosis was also increased. The exact contribution of these cell death pathways to the reduction in muscle mass observed remains to be determined. However, inhibition of ferroptosis alone enhanced cell viability following palmitate exposure while inhibition of apoptosis alone did not, possibly revealing a greater importance for ferroptosis. The role of ferroptosis in skeletal muscle pathologies is just emerging but has been implicated in the development of sarcopenia, rhabdomyolysis and inflammatory myopathies^34^. Our study also implicates skeletal muscle ferroptosis in sarcopenic obesity.

While there is much debate in the literature about the relationship between mitochondrial capacity and function and insulin action^35^, the reduction in oxidative capacity induced by HDAC4 and 5 did not result in overt insulin resistance. Overexpression of HDAC4 and 5 increased basal glucose clearance, with more glucose directed to glycogen synthesis. Oxidative utilisation of glucose was impaired through the TCA cycle, particularly at the level of succinate. The levels of succinate are thought to reflect the redox state of the mitochondrial coenzyme Q pool, with higher concentrations of reduced CoQ (CoQH_2_) preventing the oxidation of succinate by SDH^36^. Oxidation of succinate by SDH is tightly associated with reverse electron flow to complex I of the electron transport chain and reduced ROS production^37^. Indeed, succinate oxidation-dependent ROS production is critical for pro-inflammatory cytokine production by macrophages^38^, adipose tissue thermogenesis^39^ and the activity of carotid body oxygen sensing K^+^ channels^40^. In the context of the present study, HDAC4 and 5-mediated impairments in succinate oxidation and reduced ROS production likely contribute to inhibition of the ferroptosis pathway and preservation of cell viability in response to lipotoxicity. Additionally, impairments in TCA cycle activity have recently been found to promote cytosolic glutathione synthesis that increases antioxidant capacity^24^, which would also inhibit ferroptosis. Together, these data indicate that inhibition of oxidative substrate utilisation is an important adaptation to lipotoxicity.

In conclusion, this study provides evidence that in response to lipotoxicity the class IIa HDACs unite impairments in skeletal muscle oxidative metabolism with inhibition of the apoptosis and ferroptosis cell death pathways to preserve muscle integrity. These findings provide key new insights into why certain metabolic adaptations occur in response to excess lipids and advance our understanding of the aetiology of metabolic diseases driven by lipotoxicity and excess nutrient availability.

## ACKNOWLEDGEMENTS

The authors wish to thank all members of the Metabolic Reprogramming Laboratory for helpful insights and technical advice and are grateful to the technical staff of the Deakin University Animal Services team. The authors also wish to thank Hongwei Qian for AAV production. This work was supported by grants from the National Health and Medical Research Council (NHMRC: APP1027227, APP1030474 and APP1127059) to SLM. PJM, PG and MAF are supported by Fellowships from the NHMRC.

## AUTHOR CONTRIBUTIONS

Conceptualization, SDM, TC and SLM; Methodology, SDM, TC, DDS, GMK, CRB, PG, FMC, KRW and SLM; Investigation, SDM, TC, AS, KAM, DCH, BN, DDS, GMK, FMC and SLM; Resources, DLT, PJM, CRB, PG, MAF and SLM; Writing – original draft, SDM, TC and SLM; Writing – review and editing, all authors; Funding acquisition, SLM.

## AVAILABILITY OF DATA AND MATERIALS

The microarray datasets and are available at Gene Expression Omnibus (GSE215185).

