## Supplementary information for "Class IIa HDACs inhibit cell death pathways and protect muscle integrity in response to lipotoxicity"

[1]Institute for Mental and Physical Heath and Clinical Translation (IMPACT) and Metabolic Research Unit, School of Medicine, Deakin University, Geelong, Victoria 3216, Australia;
[2]College of Health and Medicine, School of Health Sciences, University of Tasmania, Launceston, Australia;
[3]Baker Heart and Diabetes Institute, Melbourne, Victoria 3004, Australia;
[4]Metabolomics Australia, The University of Melbourne, Parkville, Victoria 3010, Australia;
[5]Institute of Physical Activity and Nutrition (IPAN) and School of Exercise and Nutrition Sciences, Deakin University, Geelong, Victoria 3216, Australia;
[6]Centre for Muscle Research, Department of Anatomy and Physiology, The University of Melbourne, Parkville, VIC, Australia.
[7]Monash Institute of Pharmaceutical Sciences, Monash University, Parkville, VIC, Australia.
[8]Barwon Health, Geelong, Victoria 3220, Australia
[9]These authors contributed equally to this work
[10]Lead contact

Lead contact:

Professor Sean L. McGee
Metabolic Remodelling Laboratory
Metabolic Research Unit
School of Medicine
Deakin University, Waurn Ponds, Australia, 3216
Ph: +61 3 5227 2519


### TABLE S1: Real time RT-PCR primers

| Gene | Primer | Sequence (5’-3’) |
|---|---|---|
| *Aifm1* | Forward | CAGAGAAGAGCCATTGCCTCC |
| | Reverse | ATACAATCAGGACCCTGGCCCC |
| *Aifm2* | Forward | ACTCCTTCCACCACAATG |
| | Reverse | CGGTTCTTCAAGTCTATGC |
| *Alox15* | Forward | AGGGCTGGGGCTAATTAGGA |
| | Reverse | TGCGCAGTGAGCTAGTGAAA |
| *Apaf1* | Forward | CTTCTTTATGGTGCTGAAGATTGA |
| | Reverse | GGTGGAGTGCCTGTCTAGTGT |
| *Bax* | Forward | ATGGAGCTGCAGAGGATGAT |
| | Reverse | GAAGTTGCCATCAGCAAACA |
| *Bbc3* | Forward | AGACAAGAAGAGCAGCATCGACAC |
| | Reverse | TAGGCACCTAGTTGGGCTCCATTT |
| *Bcl2* | Forward | ACGCTCTCCACACACATGAC |
| | Reverse | GGTGGTGGAGGAACTCTTCA |
| *Bcl2l1* | Forward | TGAATGACCACCTAGAGCCTTG |
| | Reverse | CAGAACCACACCAGCCACAG |
| *Casp3* | Forward | TGGACTGTGGCATTGAGACAG |
| | Reverse | CGACCCGTCCTTTGAATTTC |
| *Casp9* | Forward | GGATGCTGTGTCAAGTTTGCC |
| | Reverse | CTTTCGCAGAAACAGCATTGG |
| *Cox7a1* | Forward | CAGCGTCATGGTCAGTCTGT |
| | Reverse | AGAAAACCGTGTGGCAGAGA |
| *Cpt1b* | Forward | TCGCAGGAGAAAACACCATGT |
| | Reverse | AACAGTGCTTGGCGGATGTG |
| *Cs* | Forward | GCCAGATCACTGTGGACATGAT |
| | Reverse | CAAGAACCGAAGTCTCATACACAAG |
| *CytC* | Forward | CATCCCTTGACATCGTGCTT |
| | Reverse | GGGTAGTCTGAGTAGCGTCGTG |
| *Echs1* | Forward | ATGGAGATGGTCCTCACTGG |
| | Reverse | CGCCATGGCTACTACGATTT |
| *Fh* | Forward | AATGACACCTTTCCCACAGC |
| | Reverse | GCTTCTGTAACCCTGGCAAC |
| *Gsl2* | Forward | CTTAGGCACTGACTACGTGCACAAG |
| | Reverse | CCCACAGATCTCCACTATATGCAGG |
| *Hadh* | Forward | ACCAAACGGAAGACATCCTG |
| | Reverse | AGCTCAGGGTCTTCTCCACA |
| *Hdac4* | Forward | CCATGAAGCACCAGCAGGAG |
| | Reverse | CTCGCCACAGCACTCTCTTTG |
| *Hdac5* | Forward | TCGCTGAGAACGGCTTTACTGG |
| | Reverse | ATGTTGGGCAGAGAAGGAGACG |
| *Idh1* | Forward | GTTGGTCTTCACCCCAAAGA |
| | Reverse | GAAATGGACTCGTCGGTGTT |
| *Mcad* | Forward | GCTAGTGGAGCACCAAGGAG |
| | Reverse | CCAGGCTGCTCTCTGGTAAC |
| *Mcl1* | Forward | AGCACATTTCTGATGCCGCCT |
| | Reverse | GTGCCTTTGTGGCCAAACACT |
| *Pmaip1* | Forward | GCTACCACCTGAGTTCGC |
| | Reverse | TCGTCCTTCAAGTCTGCTG |
| *Ppargc1a* | Forward | CCCTGCCATTGTTAAGACC |
| | Reverse | TGCTGCTGTTCCTGTTTTC |
| *Ppard* | Forward | CACTTGTTGCGGTTCTTCTTC |
| | Reverse | CCTCGGGCTTCCACTACG |
| *Sat1* | Forward | AAGTGTCGCTGCAGTAT |
| | Reverse | AGCCTCCATCCCTGTTCACT |

| | | |
|---|---|---|
| *Slc7a11* | Forward | TGGCGGTGACCTTCTCTGA |
| | Reverse | ACAAAGATCGGGACTGCTAATGA |

**TABLE S2: Antibodies**

| Antibody | Company | Dilution |
|---|---|---|
| HDAC4 | Cell Signaling | 1:1000 in TBST |
| HDAC5 | Santa Cruz | 1:1000 in TBST |
| OXPHOS cocktail | MitoSciences | 1:1000 in TBST |
| pT308 Akt | Cell Signaling | 1:1000 in TBST |
| pS473 Akt | Cell Signaling | 1:1000 in TBST |
| Akt | Cell Signaling | 1:1000 in TBST |
| Caspase 3 | Cell Signaling | 1:1000 in TBST |
| Cleaved caspase 3 | Cell Signaling | 1:1000 in TBST |
| Caspase 9 | Cell Signaling | 1:1000 in TBST |
| Cleaved caspase 9 | Cell Signaling | 1:1000 in TBST |
| acetyl K120 p53 | Abcam | 1:500 in TBST |
| p53 | Cell Signaling | 1:1000 in TBST |
| pS642 TBC1D4 | Cell Signaling | 1:1000 in TBST |
| TBC1D4 | Cell Signaling | 1:500 in 1% BSA in TBST |
| α-tubulin | Sigma-Aldrich | 1:1000 in TBST |

**SUPPLEMENTARY FIGURE 1**

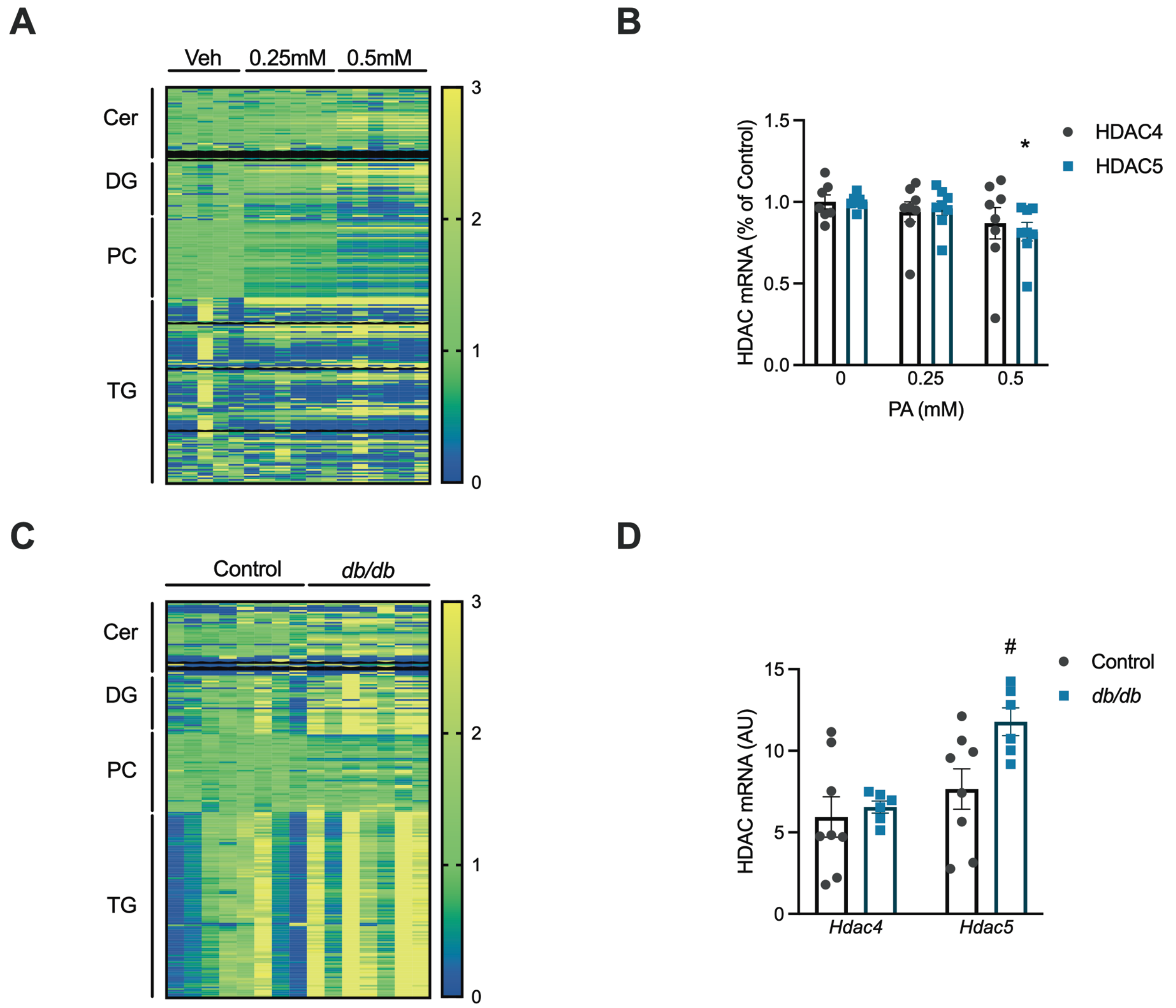


**Supplementary Figure 1:** (A) Heat map of ceramide (Cer), diglyceride (DG), phosphatidylcholine (PC) and triglyceride (TG) lipid species; (B) *Hdac4* and *Hdac5* gene expression following exposure to vehicle, 0.25mM or 0.5mM palmitate (PA) for 24hrs. (C) Heat map of Cer, DG, PC and TG lipid species; (D) *Hdac4* and *Hdac5* gene expression in tibialis anterior skeletal muscle from Control and *db/db* mice (Unpaired t-test, *p*=0.0252). Data are mean±SEM, n=6-8 biological replicates per group for cell experiments and 6-8 per group for animal experiments.

### SUPPLEMENTARY FIGURE 2

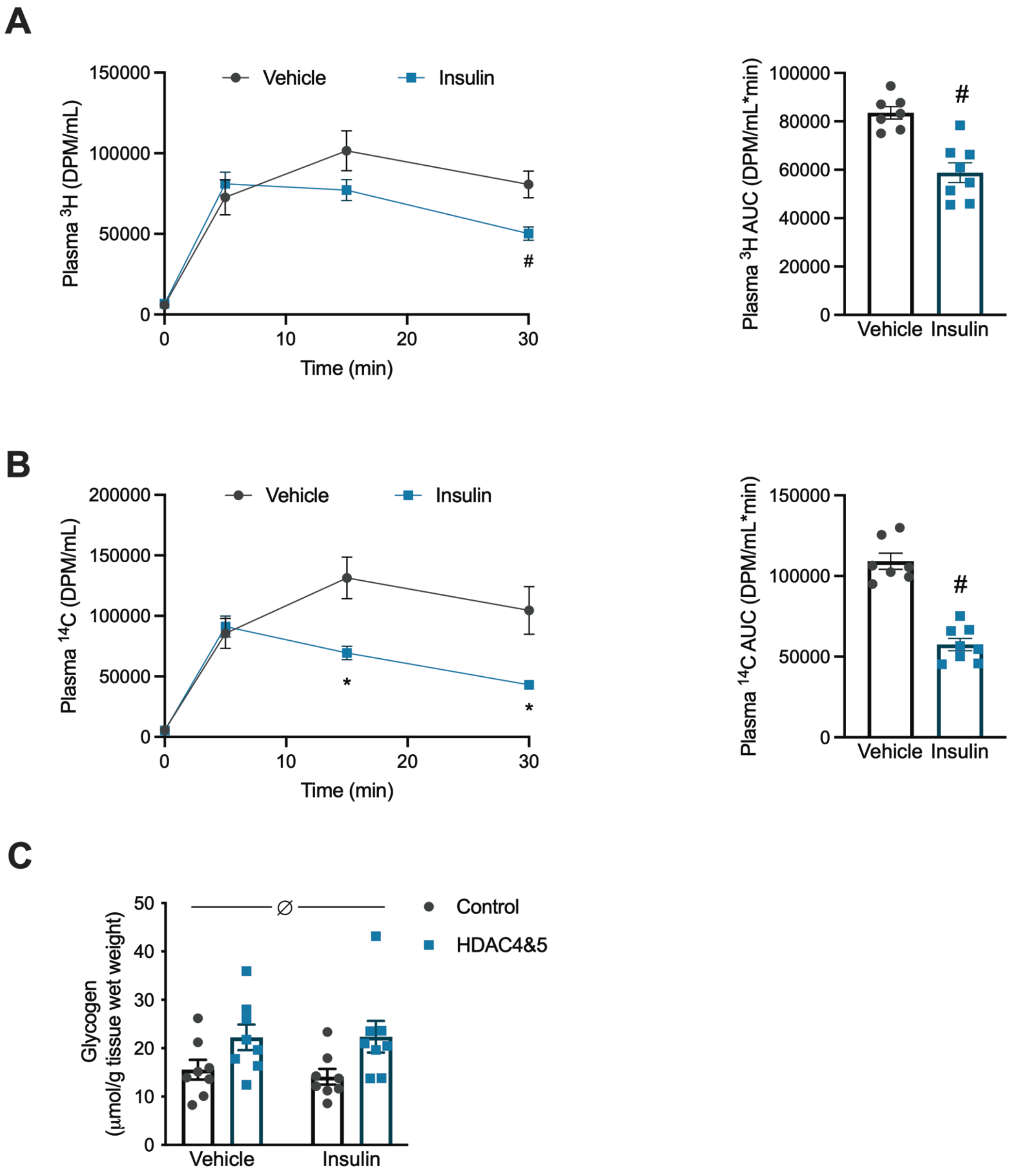


**Supplementary Figure 2:** (A) Plasma $^{3}$H at 0, 5, 15 and 30 min (Two-way ANOVA, time *p*<0.0001 *F*(2.607,36.5)=68.08, interaction *p*=0.0079 *F*(3,42)=4.511, $^{\#}p<0.05$ vs Vehicle for same time point by Tukey's multiple comparisons tests) and plasma $^{3}$H area-under-the-curve (AUC; Unpaired t-test *p*=0.0003) following vehicle or insulin and $^{3}$H-2-deoxyglucose administration. (B) Plasma $^{14}$C at 0, 5, 15 and 30 min (Two-way ANOVA, time *p*<0.0001 *F*(3,21)=47.84, treatment *p*=0.0442 *F*(1,7)=5.997, interaction *p*=0.0010 *F*(3,21)=7.944, $^{\#}p<0.05$ vs Vehicle for same time point by Tukey's multiple comparisons tests) and plasma $^{14}$C AUC (Unpaired t-test *p*<0.0001) following vehicle or insulin and $^{14}$C-glucose administration. (C) Glycogen (Two-way ANOVA, $^{\varnothing}$genotype *p*=0.0053 *F*(1,28)=9.131) in Control or HDAC4 and 5 overexpressing skeletal muscle. Data are mean±SEM, n=6-8 per group.

### SUPPLEMENTARY FIGURE 3

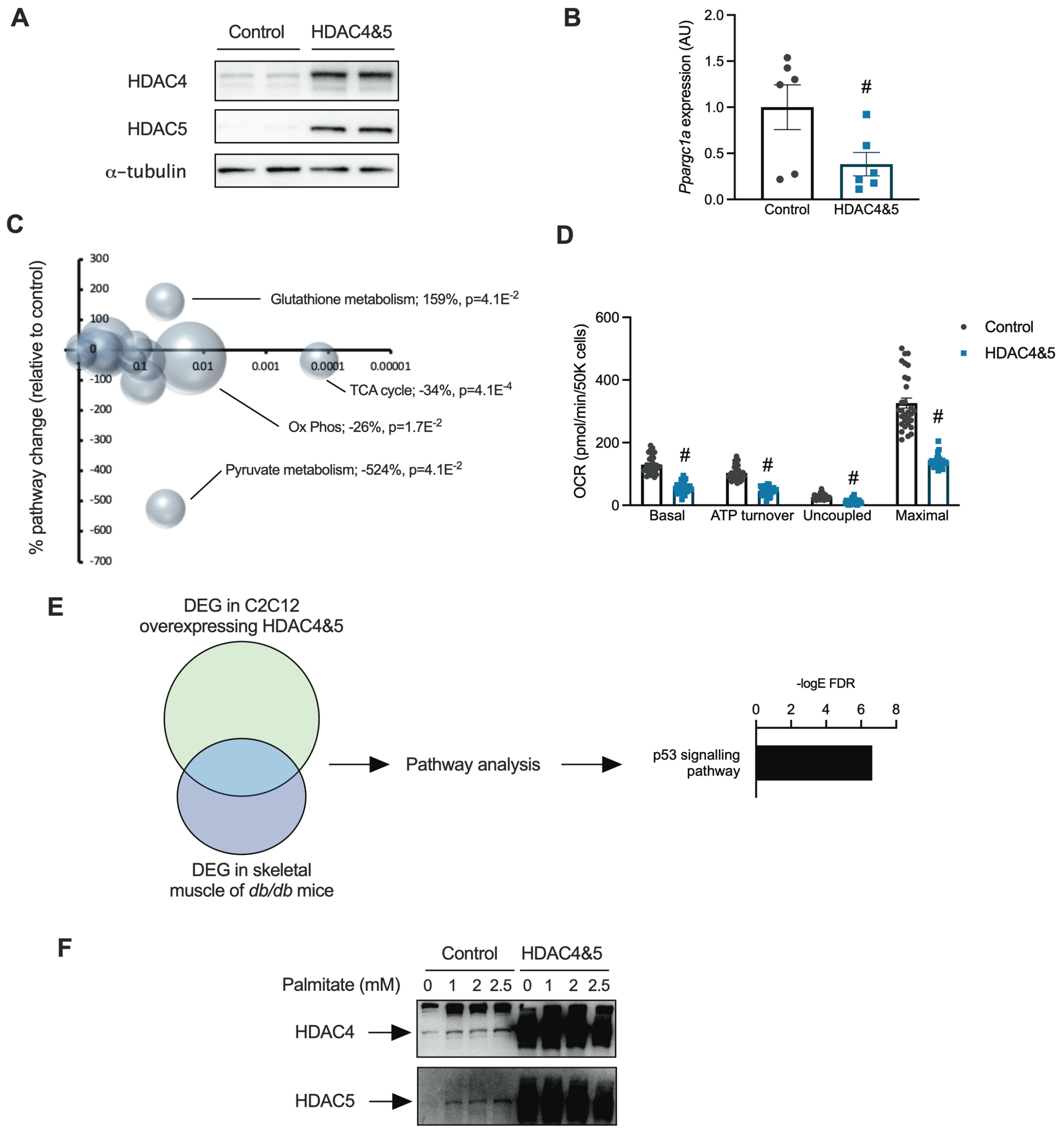


**Supplementary Figure 3:** (A) HDAC4 and 5 protein; (B) *Ppargc1a* gene expression (Unpaired t-test, $p$=0.0475); (C) Bubble plot of metabolic pathway gene expression; (D) Oxygen consumption rate (OCR) linked to basal (Unpaired t-test, $p$<0.0001), ATP turnover (Unpaired t-test, $p$<0.0001), uncoupled (Unpaired t-test, $p$<0.0001) and maximal respiration (Unpaired t-test, $p$<0.0001) in Control and HDAC4 and 5 overexpressing C2C12 myoblasts. (E) Identification of co-differentially expressed genes in skeletal muscle from *db/db* mice and C2C12 myoblasts overexpressing HDAC4 and 5 (n=3/group for skeletal muscle and cell analyses) followed by pathway enrichment analysis revealed the p53 signalling pathway as being significantly enriched in this geneset. Data are mean±SEM, n=6-20 biological replicates per group. # p<0.05 vs Control. (F) Overexposed western blots showing the increase in endogenous HDAC4 and 5 with high concentrations of palmitate.

**SUPLEMENTARY FIGURE 4**

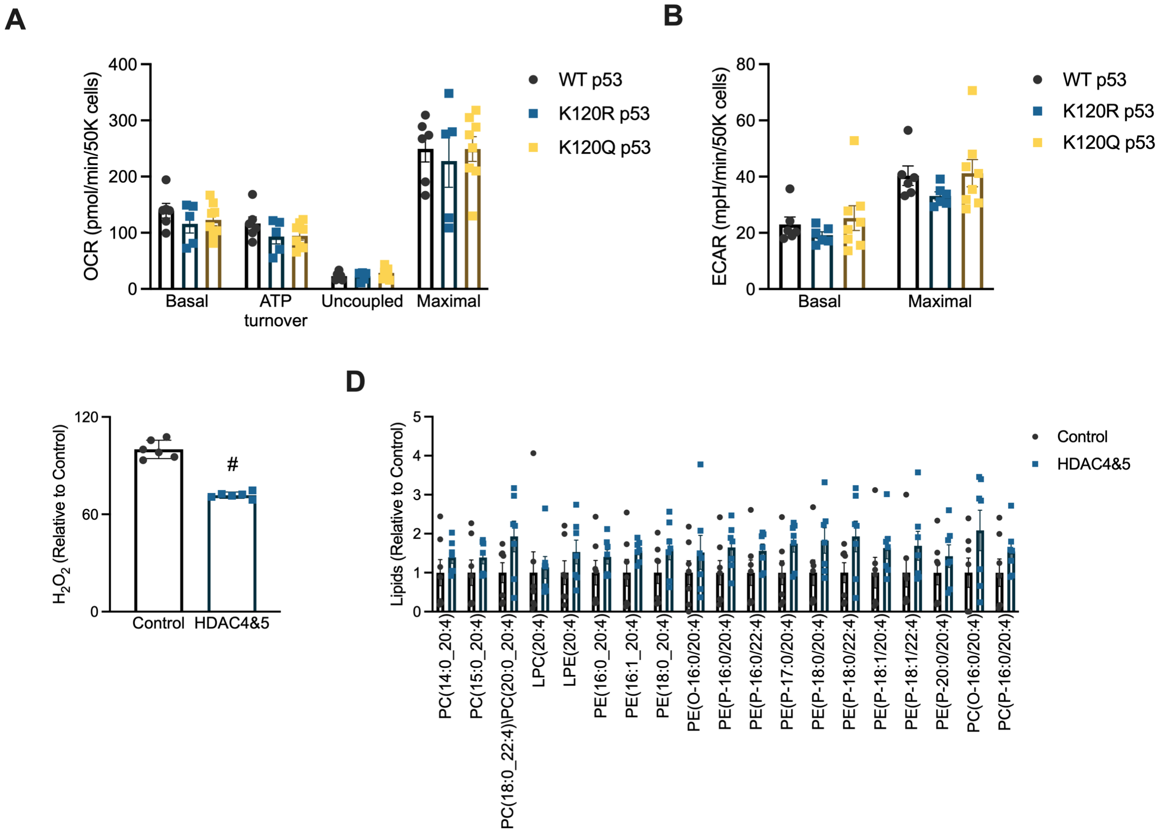


**Supplementary Figure 4:** (A) Oxygen consumption rate (OCR) linked to basal, ATP turnover, uncoupled and maximal respiration and (B) Basal and maximal extracellular acidification rate (ECAR) in C2C12 myoblasts expressing WT, K120R or K120Q p53. (C) Total $H_2O_2$ release from Control and HDAC4 and 5 overexpressing C2C12 myoblasts (Unpaired t-test, $p<0.0001$). (D) Phosphatidylcholine (PC), lysophosphatidylcholine (LPC), lysophosphatidyl-ethanolamine (LPE) and phosphatidylethanolamine (PE) lipids in Control or HDAC4 and 5 overexpressing skeletal muscle. Data are mean±SEM, n=6 biological replicates per group for cell experiments and n=7 per group in mouse experiments. # p<0.05 vs Control.

### SUPPLEMENTARY FIGURE 5

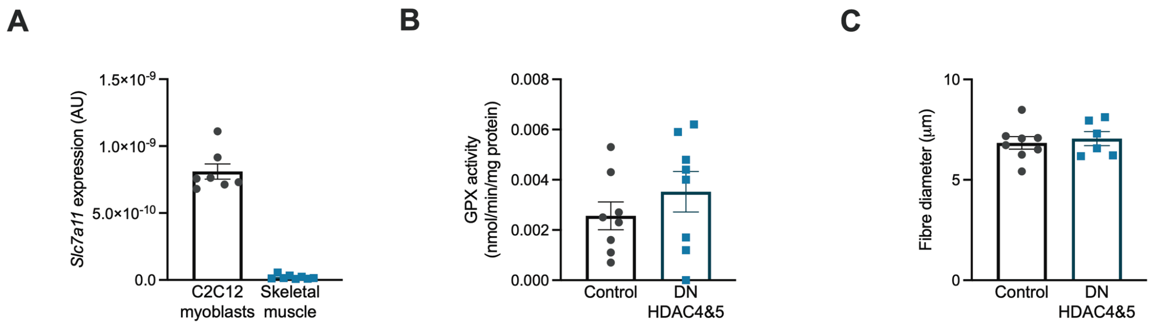


**Supplementary Figure 5:** (A) *Slc7a11* expression in C2C12 myoblasts or mouse skeletal muscle. (B) Glutathione peroxidase (GPX) activity, and; (C) Fibre diameter in Control and DN HDAC4 and 5 skeletal muscle of *db/db* mice. Data are mean±SEM, n=7-8 per group.
